# Proteomic and Genomic Characterization of a Yeast Model for Ogden Syndrome

**DOI:** 10.1101/045047

**Authors:** Max J. Döerfel, Han Fang, Jonathan Crain, Michael Klingener, Jake Weiser, Gholson J. Lyon

## Abstract

Naa10 is a Nα-terminal acetyltransferase that, in a complex with its auxiliary subunit Naa15, co-translationally acetylates the α-amino group of newly synthetized proteins as they emerge from the ribosome. Roughly 40-50% of the human proteome is acetylated by Naa10, rendering this an enzyme with one of the most broad substrate ranges known. Recently, we reported an X-linked disorder of infancy, Ogden syndrome, in two families harboring a c.109T>C (p.Ser37Pro) variant in NAA10. In the present study we performed in-depth characterization of a yeast model of Ogden syndrome. Stress tests and proteomic analyses suggest that the S37P mutation disrupts Naa10 function thereby reducing cellular fitness, possibly due to an impaired functionality of molecular chaperones, Hsp104, Hsp40 and the Hsp70 family. Microarray and RNA-seq revealed a pseudo-diploid gene expression profile in ΔNaa10 cells, likely responsible for a mating defect. In conclusion, the data presented here further support the disruptive nature of the S37P/Ogden mutation and identify affected cellular processes potentially contributing to the severe phenotype seen in Ogden syndrome.

## Introduction

Nα-terminal acetylation (NTA) is one of the most prevalent protein modifications known with 80–90% of the human proteome being acetylated at their Nα-terminus ^11–13^. NTA is catalyzed by Nα-terminal acetyltransferases (NATs) that transfer the acetyl moiety from acetyl-CoA to the α-amino group of their protein substrates. Despite its discovery more than 50 years ago, the functional consequences of this modification are not fully understood. Early studies indicated that NTA can modulate protein-protein interaction ^1–3^. This feature seems to be especially relevant for proteolytic processes where NTA increases the affinity of ubiquitin ligases towards their substrates. As such, NTA creates degradation signals (Ac/N-degrons) that can be recognized by the ubiquitin ligase Doa10 and Not4 that promote ubiquitination and proteasome-mediated degradation in conjunction with Ubc6 and Ubc7 ^4–6^. In other cases, NTA has been shown to increase protein stability by blocking the access for N-terminal ubiquitination thereby protecting proteins from proteasomal degradation ^7–10^.

Separately, there is increasing evidence that NTA can help to structurally stabilize α-helices, particularly in proteins containing an unstructured N-terminus^14–22^. Additionally, protein function has been shown to be regulated by NTA in humans^23–26^ and y east ^27–30^. Many efforts have been made to analyze the physiological role of NatA/Naa10 in the past years; however, our understanding is still incomplete. NatA function has been linked to many cellular processes, including cell cycle control, DNA-damage response, hypoxia, apoptosis and cancer. For recent reviews, the reader is referred to ^31; 32^.

To date six NATs have been discovered in humans (NatA-NatF) with a distinct substrate specificity ^33^. The best studied NAT is NatA, acetylating around 40–50% of the human and yeast proteome ^34; 35^. NatA is composed of Naa10 (Ard1p in yeast) and its auxiliary subunit Naa15 (Nat1p). Naa15 is believed to anchor the complex to the ribosome to enable co-translational NTA as soon as a newly synthetized peptide emerges from the exit tunnel ^36–38^. Bound to Naa15, the catalytic subunit Naa10 acetylates the α-amino group of proteins starting with alanine, serine, glycine, threonine, valine and cysteine, after the initiator methionine has been cleaved by methionine-aminopeptidases ^11; 39; 40^. In the monomeric form, Naa10 displays a different substrate specificity, post-translationally acetylating proteins starting with aspartate or glutamate ^12; 41^. Structural analyses showed that the reason for this shift in substrate specificities was due to conformational changes induced by the binding to Naa15 ^42^. Lysine acetyltransferase activity has also been attributed to Naa10; however, structural and biochemical data strongly suggests that Naa10 does not have the properties to facilitate direct ε-acetylation of lysine sidechains ^42; 43^.

Recently, we have identified a c.109T> (p.Ser37Pro) variant in NAA10 in two families with a lethal X-linked disorder of infancy which we named Ogden syndrome in honor of where the first family lives (in Ogden, Utah). The disorder comprises a distinct combination of an aged appearance, craniofacial anomalies, hypotonia, global developmental delays, cryptorchidism, and cardiac arrhythmias ^44^ Functional analysis of this mutation revealed a decreased enzymatic activity of the S37P mutant *in vitro* and in primary cells derived from the patients as well as impaired complex formation of Naa10 and Naa15 ^44–46^. Furthermore, we identified affected proteins which were less NTA in patient cells, one of which is THOC7 (THO complex subunit 7 homologue). Notably, the reduced acetylation of THOC7 was associated with a decreased stability of the protein, possibly contributing to Ogden syndrome ^45^.

Since the first description of Ogden syndrome in two multiplex unrelated families, additional variants in Naa10 have been identified in singleton probands with non-syndromic intellectual disabilities accompanied with postnatal growth failure and skeletal anomalies ^47; 48^,one family with two brothers with syndromic intellectual disability with long QT ^49^ and one multiplex family with Lenz microphthalmia syndrome (LMS), characterized by microphthalmia or anophthalmia, developmental delay, intellectual disability, skeletal abnormalities and malformations of teeth, fingers and toes ^50^. The phenotypic differences between all cases are quite distinct, and to date there has been no unifying explanation for this.

Here, we expanded on a prior study^51^ by using *S. cerevisiae* to study the impact of Naa10 disruption in several different physiologic situations and by conducting genomic and proteomic assays with an emphasis on the S37P/Ogden mutation.

## Materials & Methods

### Yeast strains

Derivatives of parental stain W303–1A (*leu2–3,112 trp1–1 canl–100 ura3–1 ade2–1 his3–11,15*) were used to generate strains with variations of the endogenous *yNAA10* and *yNAA15* loci. To introduce the human S37P mutation in yeast (YG36), first the homolog position was identified as Serine 39 by sequence alignment (Figure 1A). *yNAA10* was amplified from W303–1A using the primers 5'-GTA GAA TTC GCC GCC ATG CCT ATT AAT ATT CGC AG and 5'-CAT GAA TTC CCT ACC GAA TTA GCA CTG CAG T and cloned into pBEVY-U (Addgene stock #51230). The yS39P mutation was introduced using the QuikChange Multi Site-Directed Mutagenesis Kit (Agilent) and the 5′-ATG TAT CAT ATT CTC CCG TGG CCG GAG GCT T primer. *yNAA10–S39P* was then PCR amplified using 5'-GGG AAA CCT AAA TAC ATA CGA TCA AGC TCC AAA ATA AAA CTT CGT CAA CCA TGC CTA TTA ATA TTC GCA GAG CG and 5'-TGT GAA GAA GCC TGG ATG AAA ATA TAC TAC GTT TAT ATA GGT TGA TTT AAT TAT ACA ATG ATA TCA TTT ACG CCT TGC primers and the resulting PCR product was transformed into the deletion stain YG10 (ΔyNAA10::URA3–15) using the lithium acetate technique ^52^. Transformants were selected using 5–FOA and verified by sequencing of the *yNAA10* locus followed by Western blot analysis (Figure 1B). The double deletion strain (YG67: ANaa10/ANaa15) was generated by crossing YG3 and YG10 to form a diploid cell (grown on SC^-Leu,Ura^). Once the cells diploidized, tetrad dissections were performed to isolate a haploid double knockout. Candidates were selected on SC'^Leu,Ura^ plates and screened for mating type and auxotrophies by replica plating.

**Figure 1:**
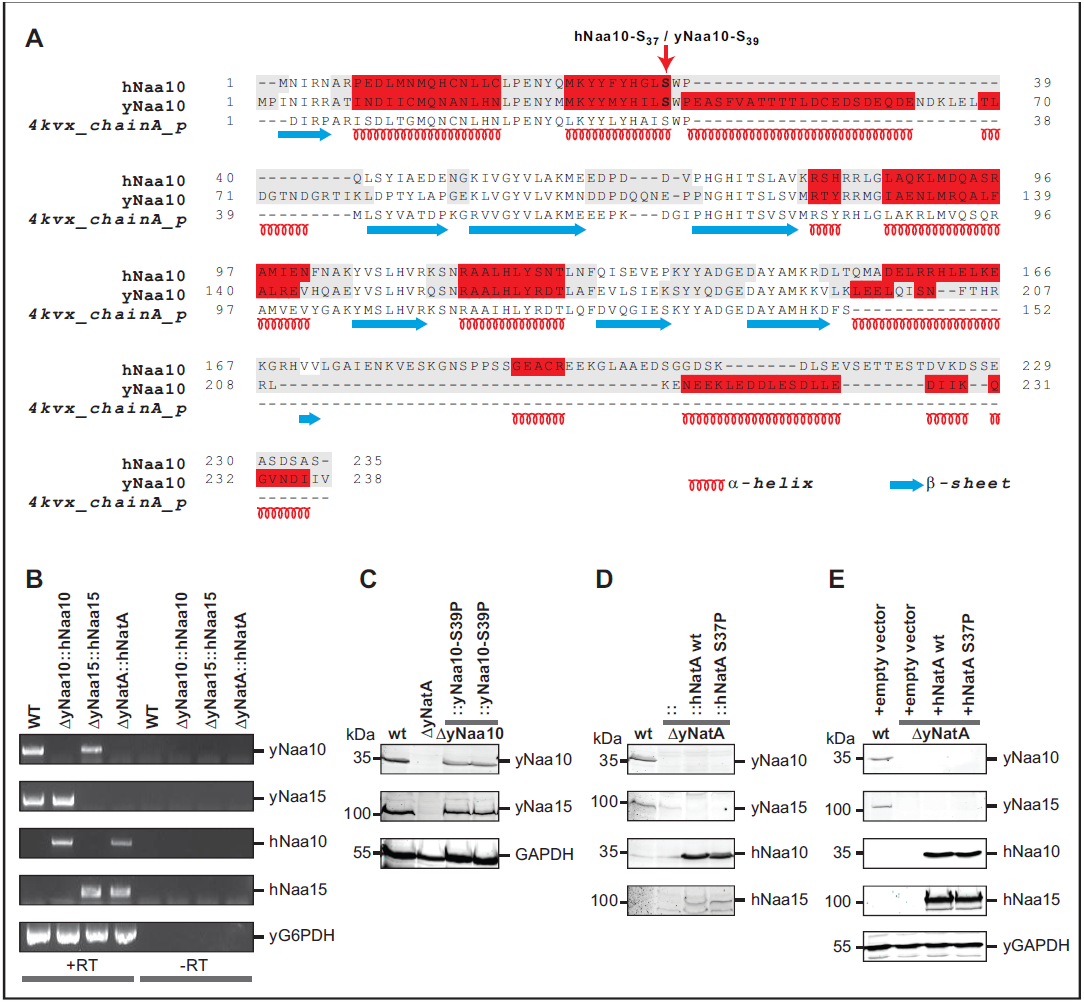
Characterization of strains used in this study. A) Sequence alignment of human and *S. cerevisiae* Naa10. Protein sequences were aligned using PROMALS3D ^82^ and the crystal structure of Naa10 from *S. pombe* (PDB structure 4KVX) ^42^ was used for secondary structure prediction. The alignment matrix between human and yeast Naa10 reached a 49.5 % identity score. The homologous position to hNaa10 S37P (yeast Serine 39) was marked with an arrow. B) RT-PCR characterization of the human replacement strains using primers designed to amplify regions internal to the indicated transcripts. The products of these reactions are shown, in the presence and absence of reverse transcriptase. C) Western blot analyses with antibodies for the human and yeast Naa10 and Naa15 along with yGAPDH as a loading control for the strains where the yNaa10 has been modified, D) for the humanized strains, expressing the human proteins from the endogenous locus and E) for the strains overexpressing the human proteins from plasmids.

To “humanize” yeast, the *yNAA10* and *yNAA15* loci were replaced by the corresponding human cDNA. First, hNaa10 WT or S37P was amplified from pBEVY-U-hNAA15-hNAA10 (Arnesen et al., 2009) with primers adding homology to the *yNAA10* locus (Supplemental Table 3:ON41/ON42: 5'-GGG AAA CCT AAA TAC ATA CGA TCA AGC TCC AAA ATA AAA CTT CGT CAA CCA TGA ACA TCC GCA ATG CGA GGC C and 5'-TGT GAA GAA GCC TGG ATG AAA ATA TAC TAC GTT TAT ATA GGT TGA TTT AAT CAG GAG GCT GAG TCG GAG GCC). Similarly as above, the PCR product was transformed into strain YG10 (yNaa10Δ::URA3–15) and transformants were selected using 5-FOA. To replace the *yNAA15*, hNaa15 cDNA was amplified from pBEVY-U-hNAA15-hNAA10 using ON54/ON55 (5'-CGT ACG ATA TCA TGC CGG CCG TGA GCC TCC CGC CCA A and 5'-GGC TAG ATA TCT CAA ATT TCA TTG GCC AGT TCT TCA GC) and cloned into pUG6-tTA ^53^ to generate an hNaa15-KanMX cassette. The hNaa15–KanMX cassette was then amplified using ON59/ON59 (5'-GGA AGG CGA TTG ACC CTA ACG AAG TAT GCC GGC CGT GAG CCT C and 5'-AAT TGA CAC ATT GAG GAG TTG CAG GAA GCT CCT CGA GCG TCG ACA) to add homology for the *yNAA15* locus. Additional homology was added using ON60/ON61 (5’ CCT TGT TCA AGA CAA ATA CCA TTG AGG AAG GCG ATT GAC CCT AAC and 5’ TAT ATA CAT AAA TTA AGT AAG AGT TAA TTG ACA CAT TGA GGA GTT). The PCR product was transformed into the respective yNaa10:hNaa10 strain and transformants were identified by selection on kanamycin plates. All strains generated in this study were validated by PCR and/or Sanger sequencing. For PCR verification, primers were designed to bind outside the intended sites of recombination such that a slower migrating product would be observed in the case of a correctly targeted transformation (Supplemental Table 3: ON87/ON48). For Sanger sequencing verification, we confirmed that the genomic region immediately adjacent to the inserted transformation product corresponded to the SGD W303 reference sequence of the intended insert site. Further, we observed the absence of yArd1 and yNat1 transcripts in the humanized strains (yNatA::hNatA WT and yNatA::hNatA S37P) by RT-PCR (Figure 1B). The indicated *S. cerevisiae* strains were harvested during mid-exponential phase growth in YPD, and total RNA was extracted using the RNeasy Midi Kit (Qiagen). Following RNA extraction, mRNA was reverse transcribed to cDNA using Superscript III and a poly(T) primer as per the manufacturer's protocol. The resulting cDNA was amplified using PCR with primers designed to hybridize to regions internal to the indicated transcripts (Supplemental Table 3: yON105–114). To control for the presence of genomic DNA, PCR reactions were run in parallel with an aliquot of the reverse transcription reaction that was run in the absence of Superscript (Figure 1B, right side). In addition, expression of human and yeast Naa10 and Naa15 in the respective strains was verified by Western blot using antibodies specific to human and yeast Naa10 and Naa15 (see Figure 1C–E). A list of all strains used is shown in Table 1.

**Table 1:**
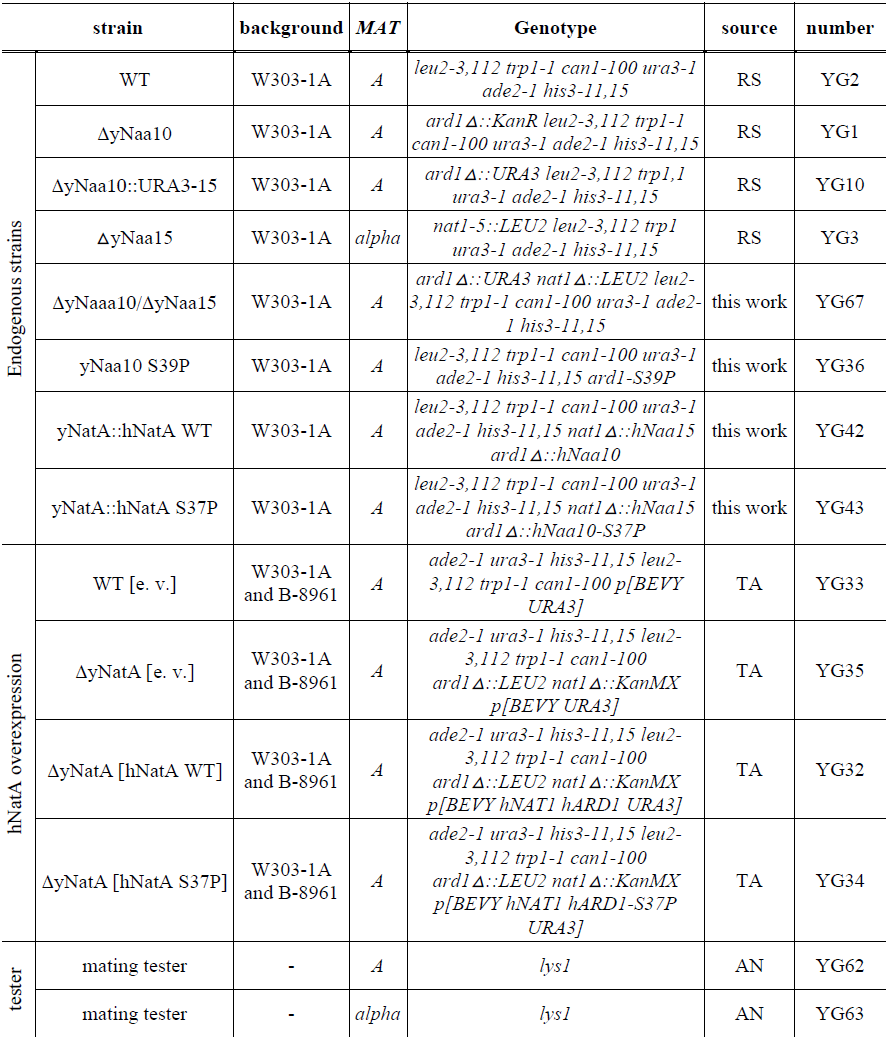
Yeast strains used in this work. Abbreviations:RS, Rolf Sternglanz; TA, Thomas Arnesen; AN, Aaron Neiman

### Growth assay in liquid culture

>A single colony was picked and grown in 5 ml YPDA media (Clontech; endogenous strains) or SC^-URA^ (Clontech, overexpressing strains) at 30°C overnight. The culture was diluted, grown for 2 further doublings, diluted to an OD_600_ of 0.1 and incubated at 30°C or 39° for 24 h. The OD_600_ was monitored and OD plotted for time point 24 h. The experiment was performed at least 7 times.

### Stress test on solid media

Similar to the stress tests in liquid culture, a single colony was grown in 5 ml of the corresponding media overnight, then diluted to an OD_600_ of 0.1. Subsequently, serial 1:5 dilutions were prepared and 10 μl of these spotted manually with a pipettor on YPDA or SC^-URA^ plates. All plates were incubated for 24–48 h at 30°C or 39°C to induce heat shock. For the chemical treatments, cells were spotted on plates containing 0–0.8 μg/ml cycloheximide or 0–400 μg/ml doxorubicin. For the UV treatment, plates were irradiated with 0, 15, and >20 mJoule in a UV Stratalinker 2400 (Stratagene) after the cells were spotted. All experiments were performed at least 3 times.

### Western Blot

To analyze protein levels in yeast, a 5 ml culture was grown in exponential phase, the yeast pelleted at 2.000 × g for 5 min and washed in H2O. Cell pellets were resuspended in 100 μl 0.2 M NaOH, incubated for 5 min at room temperature, mixed with 50 μl lysis buffer (65 mM Tris, pH 7.5; 4 % (m/v) SDS, 4 % (v/v) β-mercaptoethanol) and incubated at 96°C for 5 min. The protein concentration was determined using APA assay (Cytoskeleton) and equal amounts were separated by SDS PAGE. The following primary antibodies were used: hNaa10 (Protein Tech, #14803–1– AP), hNaa15 (Abcam, #ab60065), yGAPDH (Sigma, #A9521–1VL), and yNaa10 and yNaa15 (kindly provided by Sabine Rospert). Anti-mouse IRDye 800CW and anti-rabbit IRDye 680RD (LI-COR) were used as secondary antibodies.

### Mass spectrometry

For protein expression analyses yeast were harvested during the exponential phase of growth at an OD_600_ of 0.6, separated by filtration (see below) and cryogenically grinded in a SPEX Freezer Mill. The samples were resuspended in Y-PER yeast protein extraction reagent (Thermo Fisher) and cellular debris removed by centrifugation. Each sample was chemically labeled separately with 1 of up to 8 distinct isobaric iTRAQ reagents^54^. Peak lists were extracted using the MASCOT Distiller software (MatrixScience). False discovery rates (FDR) were estimated by searching equivalent reversed or randomized sequence databases, using a cutoff value of 1% FDR. Protein-level expression ratios were calculated using intensity-weighted averages from matching peptides after outlier removal. This experiment was performed in duplicate. Comparing the two sample replicates, we fitted a Cauchy distribution using the EasyFit statistical package (MathWave, Inc.). Peptide p values were calculated using a two-tailed t-test.

### Microarray

For gene expression analysis, RNA was extracted from a 10 ml yeast culture growing at exponential phase (OD_600_ of 0.6) at 30°C using phenol-chloroform extraction method. Briefly, cells were harvested by filtration through a 25 mm Nylon Hydrophilic Membrane Filter (Whatman), resuspended in 750 μl lysis buffer (10 mM EDTA; 0.5 % SDS; 10 mM Tris, pH 7.5), mixed with an equal volume of saturated phenol, pH 4.3 (Fisher Scientific) and incubated at 65°C for 60 min with vortexing every 20 min. The samples were incubated on ice for 10 min, centrifugated for 5 min at 5,000 g and the supernatant was transferred to 2 ml phase lock gel tubes (PLG) heavy (VWR) and mixed with 750 μl chloroform. The aqueous phase was collected according to the manufacturer’s instructions. The RNA was precipitated with 75 μl 3 M sodium acetate and 1.5 ml ethanol for 60 min at –20°C, pelleted at 3,000 rpm for 10 min, washed twice with 70% ethanol and air dried at room temperature for 30 min. The RNA was dissolved in 25 μl H2O at room temperature, further purified using a RNeasy Mini Kit (Qiagen) and labelled with a Quick Amp Labeling Kit (Agilent Technologies) using cyanine–3–CTP/cyanine-5–CTP (Fisher Scientific). The probes were hybridized to 8x15k yeast microarrays and scanned using a 2–μm Agilent microarray scanner. The images were analyzed using the feature extraction software and normalized by global lowess using Goulphar. For each gene, the Cy5/Cy3 ratios corresponding to the duplicated probes were averaged. The averaged log2 ratios and the standard deviation between the two replicates were calculated for each gene.

### RNA-Seq

Total RNA was purified according to the RiboZero Gold Kit (Epicentre) and the RNA Clean and Concentration Kit (Zymo Resaearch). Libraries were generated, PCR amplified, purified using the Agencourt AMPure XP system (Beckman Coulter) and characterized on High sensitivity DNA assay (Agilent). The libraries were sequenced on the Illumina NextSeq platform under high output mode, resulting in 76 single-end reads. For quality control, we used FastQC (v0.11.3) to generate diagnostic statistics of the data and used Fastx (version 0.1) to remove low-quality reads. Then we used cutadapt (v1.7.1) to identify, remove 3’ adapter sequence (AGATCGGAAGAGCACACGTCT) from the reads, and clip the first base from the 5’ end
(options:-O 6-m 25-n 1-e 0.15-cut 1). Bowtie (v1.1–1) was used to identify and remove rRNA reads (option:-seedlen=23). We used STAR (v2.4.0j) to align the RNA-Seq reads to the Saccharomyces cerevisiae reference genome S288C (release R64–2–1) (options for STAR: --outSAMstrandField intronMotif --outSAMunmapped Within --outFilterType BySJout --outFilterMultimapNmax 20 --alignSJoverhangMin 8 --alignSJDBoverhangMin 1 --outFilterMismatchNmax 999 --outFilterMismatchNoverLmax 0.04 --alignIntronMin 0 --alignIntronMax 5000 --alignMatesGapMax 5000 --outSAMattributes NH HI AS NM MD --outSAMtype BAM SortedByCoordinate-outFilterIntronMotifs RemoveNoncanonical). Cufflinks (v2.2.1) was used to quantify RNA abundance at the gene level. We used DESeq2 to perform differential expression analysis, comparing the following combinations: k/o vs. wt, hNAA10 vs. wt, and hNAA10S37P vs. wt. Only genes with Benjamini & Hochberg (BH) adjusted p-values less than 0.05 and log2 fold changes >1 or <–1 are considered significantly differentially expressed. Telomeric regions are defined by taking the distal 40 Kb regions of either end of the chromosomes.

### Mating assay

The mating efficiency was assessed in a quantitative mating assay. Mating tester cells and cells of interest were grown in log phase to OD_600_ of 0.25. To generate tester plates, 0.1 ml of the tester strain were seeded on SC^-Lys,—Ura^ selective plates and dried for 20 min. For each strain of interest, four 1:10 dilutions were prepared and 0.1 ml seeded on top of the tester plates. To determine the initial seeding number, 70 μl were seeded on SC^-Ura^ selective plates. Cell colonies were counted after 2–3 days incubation at 30°C and the mating efficiency determined as the ratio between the numbers of colonies grown on the tester plates and initial number of seeded cells. The experiment was performed 9 times.

## Results

We previously identified a disease, Ogden syndrome, associated with a S37P variant in the human Nα-terminal acetyltransferase, Naa10^44^. To study the impact of this mutation *in vivo*, we have studied different *S. cerevisiae* Naa10 variant stains and characterized the consequences induced by this mutation under various stress conditions.

### The S37P mutation renders Naa10 functionally impaired

It was previously shown that disruption of Naa10 or Naa15 increases the sensitivity of yeast cells toward heat shock ^55–57^ To test whether the Ogden mutation functionally impairs Naa10 in yeast, we performed growth experiments at an elevated temperature. In line with previous results, all tested knockout strains showed reduced growth at 39°C (see Figure 2). The replacement of the yeast NatA complex with the human genes on the endogenous locus did not rescue the observed effects seen in the knockout (p ;= 0.195 for yNatA::hNatA WT and p = 0.595 for yNatA::hNatA S37P) and already exhibited a slight but statistically significant (p < 0.01) growth defect at 30°C (Figure 2A). Overexpression of the human NatA complex on the other hand complemented the strong defect to WT levels, whereas overexpressing the mutated hNaa10 S37P/hNaa15 only partially restored the growth defect (Figure 2B). To test whether the Ogden mutation has a similar effect on the function of the yeast Naa10, we first identified the homologous position using sequence alignment (Figure 1A) of human and *S. cerevisiae* Naa10. Mutation of this position (yS39P) did not result in a heat shock phenotype (Figure 2A).

**Figure 2:**
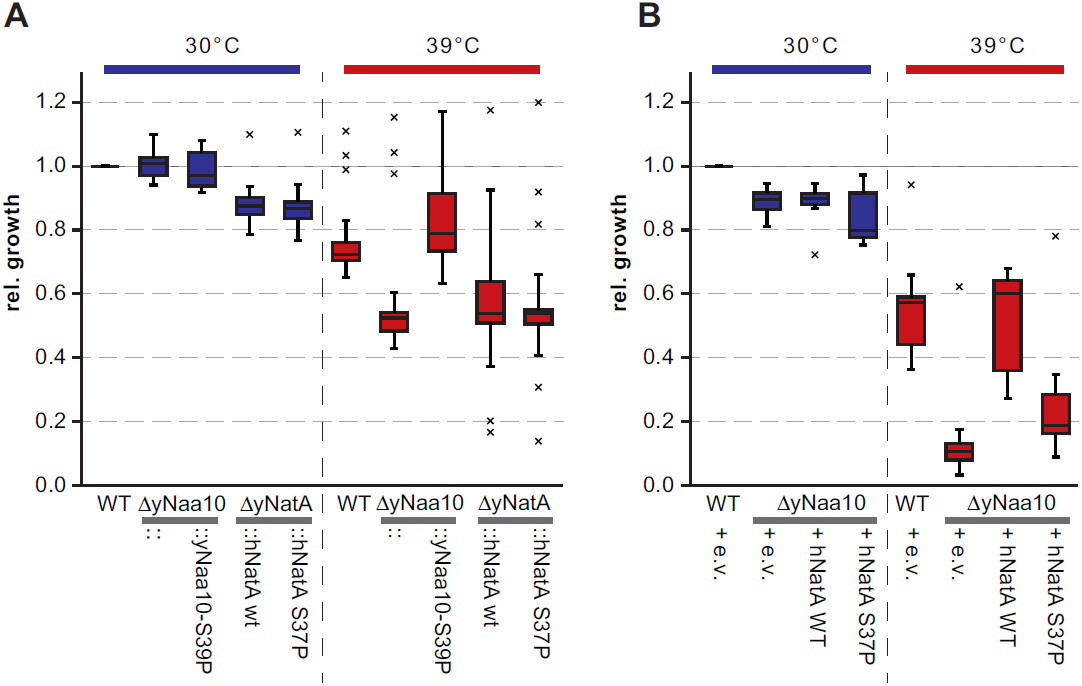
Heat stress in liquid culture. 5 ml yeast cultures were grown at 30°C or 39°C and optical density was monitored. Shown is a box plot of the relative growth after 24 h for at least 7 independent experiments for each strain under each condition. Values outside 1.5 times the interquartile range were considered outliers (and marked as such with an x), and statistical analyses were performed using student’s t-test. A) shows the results for the endogenous strains, B) shows the overexpressing strains. e.v. = empty vector

To confirm these results, stress tests on plates were performed. Similarly as for the experiments in liquid culture, the knockout of the yeast NatA complex strongly reduced the ability of the cells to grow at elevated temperatures and this effect could not be rescued by replacing the yeast loci with the human NatA genes (Figure 3A). Overexpressing the human NatA proteins on the other hand partially rescued the heat shock, with the mutant S37P being less efficient (Figure 4A).

**Figure 3:**
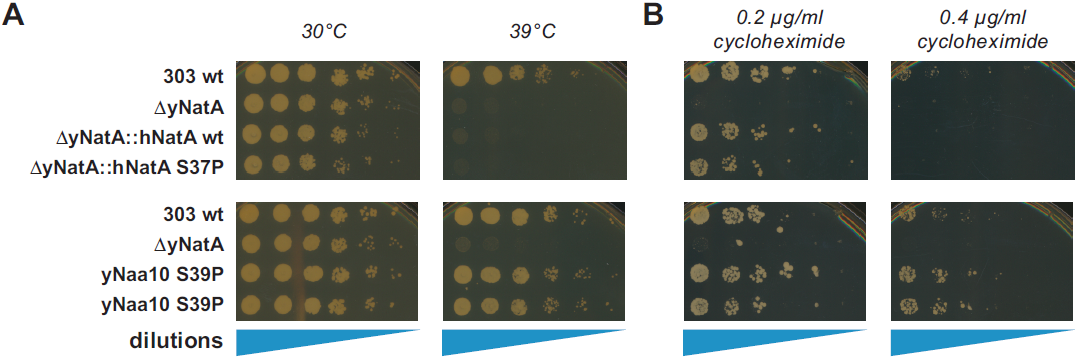
Heat and cycloheximide stress tests for endogenous strains during growth on solid media. Serial dilutions of the indicated yeast cultures were spotted on plates and incubated at 30 or 39°C (A). Cells were grown on plates containing the indicated concentrations of cycloheximide (B). All experiments were done at least in triplicate; shown are representative scans of the plates.

**Figure 4:**
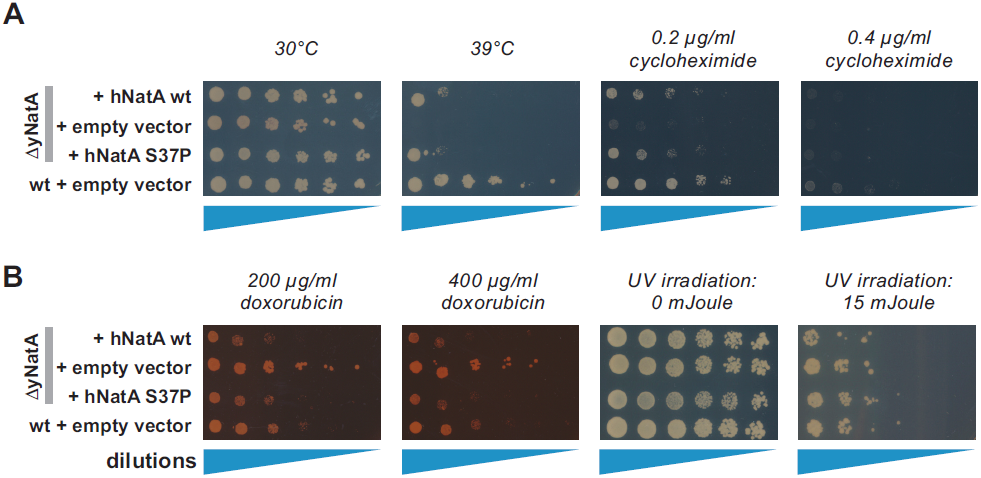
Yeast spotting tests for overexpressing strains. The yeast strains were diluted and spotted on plates and incubated and either subjected to heat shock A), cycloheximide A), doxorubicin or UV radiation stress B). All experiments were done at least in triplicate; shown are representative scans of the plates.

In addition to the heat stress, cycloheximide, doxorubicin and UV irradiation were used as stress factors. Cycloheximide blocks the translocation step in elongation and was used to challenge protein biosynthesis. Doxorubicin is an agent that intercalates into DNA, thereby trapping topoisomerase II covalently bound to DNA which subsequently induces double-strand breaks ^58^. Similarly, elevated doses of UV radiation induce DNA-damage response pathways. Both were used since Naa10 has been previously connected to the regulation of cell survival in response to DNA-damage [for a recent review see ^32^].

In analogy to the heat stress experiments, knockout of yNatA strongly affected growth on cycloheximide plates (Figure 3B and Figure 4A). This growth defect could partially be rescued by overexpressing the human hNat15/hNaa10 complex from plasmids and-in contrast to the heat shock-even by expressing the human proteins from the endogenous yeast locus (Figure 3B). Interestingly, the homologue yS39P mutation of the yeast Naa10 showed a similar growth pattern as the WT strain.

In the challenge with doxorubicin, knockout of yNatA clearly increased survival toward doxorubicin stress whereas overexpression of the human NatA complex or the human S37P mutant complex seemed to decrease growth compared to WT (Figure 4B). No obvious differences in the phenotype could be observed between strains after UV-treatment (Figure 4B); at doses ≥ 20 mJoule none of the strains survived (not shown).

### Deletion of NatA is associated with a dysregulation of heat shock proteins

As discussed above, NTA regulates specific protein function, stability and expression. Therefore, we asked whether the phenotypic effects observed might be related to changes in the abundance of one or several proteins. To analyze the expression changes induced by the knockout or overexpression of NatA, we performed quantitative mass spectrometric analyses. In total, 3191 proteins were identified from which 491 were significantly dysregulated (fold changes > 1.37 or < 0.65 significant to > 2 SD; p=0.05) (Supplemental Table 1). A GO-term enrichment analyses using WebGestalt ^59; 60^ of proteins with significantly elevated protein levels in the knockout strain (271 proteins) did not give any obviously interpretable results, except biosynthetic and metabolic processes (not shown). Proteins that were significantly less abundant in the knockout (220 proteins) were enriched for energy metabolism processes such as “ATP synthesis coupled electron transport” (GO:0042773, adjP=0.0019) or “cellular respiration” (GO:0045333, adjP=0.0033) and mating-related processes (“adaptation of signaling pathway by response to pheromone involved in conjugation with cellular fusion”, GO:0000754, adjP=0.0180). Furthermore, proteins that were dysregulated in the knockout by at least 25% but whose expression could be rescued by expression of the human NatA to within 10 % of the WT level (172 proteins), were enriched for protein folding [GO:0006457, adjP=0.0191] and protein refolding [GO:0042026, p = 0 .0705]. Protein folding and re-folding is accomplished by molecular chaperones, and from studies with Sup35/[PSI+] prion strains, it is known that loss of NatA increases heat-shock response (HSR) and increases Hsp104, Ssa1/2, Ssb1/2 and Sis1 protein levels ^61^. Interestingly, in the analysis performed here, knockout of NatA was associated with a strong increase in the Hsp70 family proteins Ssa1-4 as well as Hsp104 and Sis1, and their expression was restored upon overexpression of hNatA WT (Figure 5)

**Figure 5:**
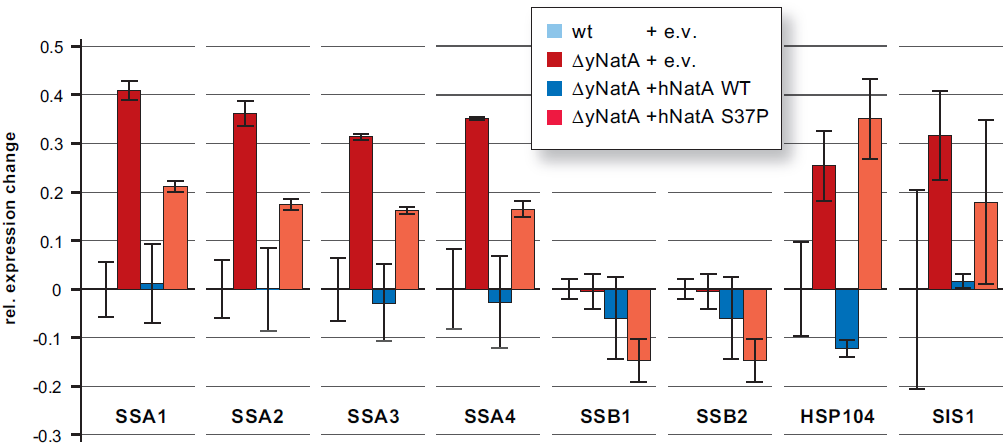
Mass spectrometric analyses of the overexpressing strains. Cells growing at exponential phase were harvested, lysed and the clarified lysates were subjected to quantitative mass spectrometry. The geometric mean of the measured abundances (relative expression) and geometric standard deviation for each protein were calculated from two independent experiments and normalized. Shown is the difference of the expression compared to the WT strain for the HSP70 chaperones Ssa1–4, Ssb1–2 as well as HSP104 and Sis1.

The Hsp70 proteins Ssb1 and Ssb2 were not found to be changed. Overexpression of the hNatA S37P mutant failed to restore the expression of the Ssa1–4, Hsp104 and Sis1 proteins. Additionally, Ssb1 and 2 were slightly less abundant in this over-expressing strain, as compared to the WT strain.

### The knockout of Naa10/NatA induces a pseudo-diploid gene expression

We next questioned whether the Naa10 knockout and the Ogden mutation might have an effect on gene expression. To address this, we performed gene expression analyses using microarray and RNA-Seq. Three different sets of yeast strains were used in the microarray study that were analyzed jointly to capture biological variability: Set 1 included a W303 WT strain, a yNaa10 knockout strain and a strain where the endogenous locus of the yeast genes were replaced with human *NAA15* and *NAA10* WT or *NAA10* S37P, respectively (see Tables

Table 1': strains YG2, YG1, YG42 and YG43). Set 2 is identical to Set 1 except instead of the single yNaa10 knockout a double yNaa10/yNaa15 knockout was used (strains YG2, YG67, YG42 and YG43). Set 3 consisted of the overexpressing strains (strains YG33, YG35, YG32 and YG34). For RNA-seq, two biological replicates of the overexpressing strains were used (YG33, YG35, YG32 and YG34).

Figure 6A shows the gene expression levels for the analyzed NATs from the microarray study, confirming the knockout of Naa10 and/or Naa15 in the corresponding strains. Also, the data shows that the deletions did not affect the expression levels of the other NATs. A table of the differentially expressed genes from the RNA-seq experiment is shown in Supplemental Table 2. In Figure 7A a Top 10 list of the genes is shown that are down-or upregulated in all knockout strains. This also revealed that only the overexpression of the human proteins from plasmids could partially rescue the effects seen in the knockout whereas the expression from the endogenous locus did not. To quantify this effect, we looked at the genes there were consistently deregulated in the knockout of all 3 sets [log2(FC) <-1 or log2(FC) >1, 39 genes]. Then we calculated the average fold expression change for the up-and downregulated genes in each condition for the endogenous sets and the overexpression set. These genes were less de-regulated upon overexpression of the human NatA complex illustrated by a reduction of the average deviation from the WT expression levels (Figure 6B). In contrast to this, when the human complex was expressed from the endogenous locus, the average expression change compared to WT was similar as seen upon knockout of yNatA, indicating no rescue effect.

**Figure 6:**
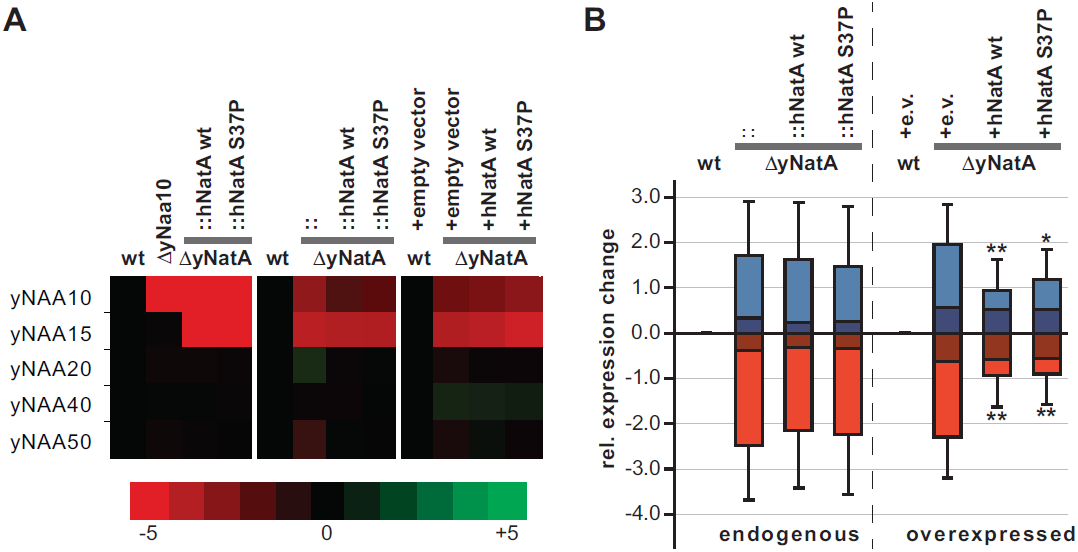
Overexpression of hNatA complements the changes induced by disruption of yNatA. A) Heat map representation of the microarray results for the NATs. B) Shown is the average expression change (and standard deviation) of up-(blue) and down-regulated (red) genes for the endogenous rescue strains (left) and the overexpression strains (right). The darker shades show the results for when all genes were plotted, whereas the lighter shades show only those genes whose expression was changed by at least ±1 in all knockout strains. Wilcoxon rank sum test was used to analyze whether the expression change was significantly reduced in the rescue strain when compared to the corresponding knockout. ** p < 0.001; * p < 0.05

**Figure 7:**
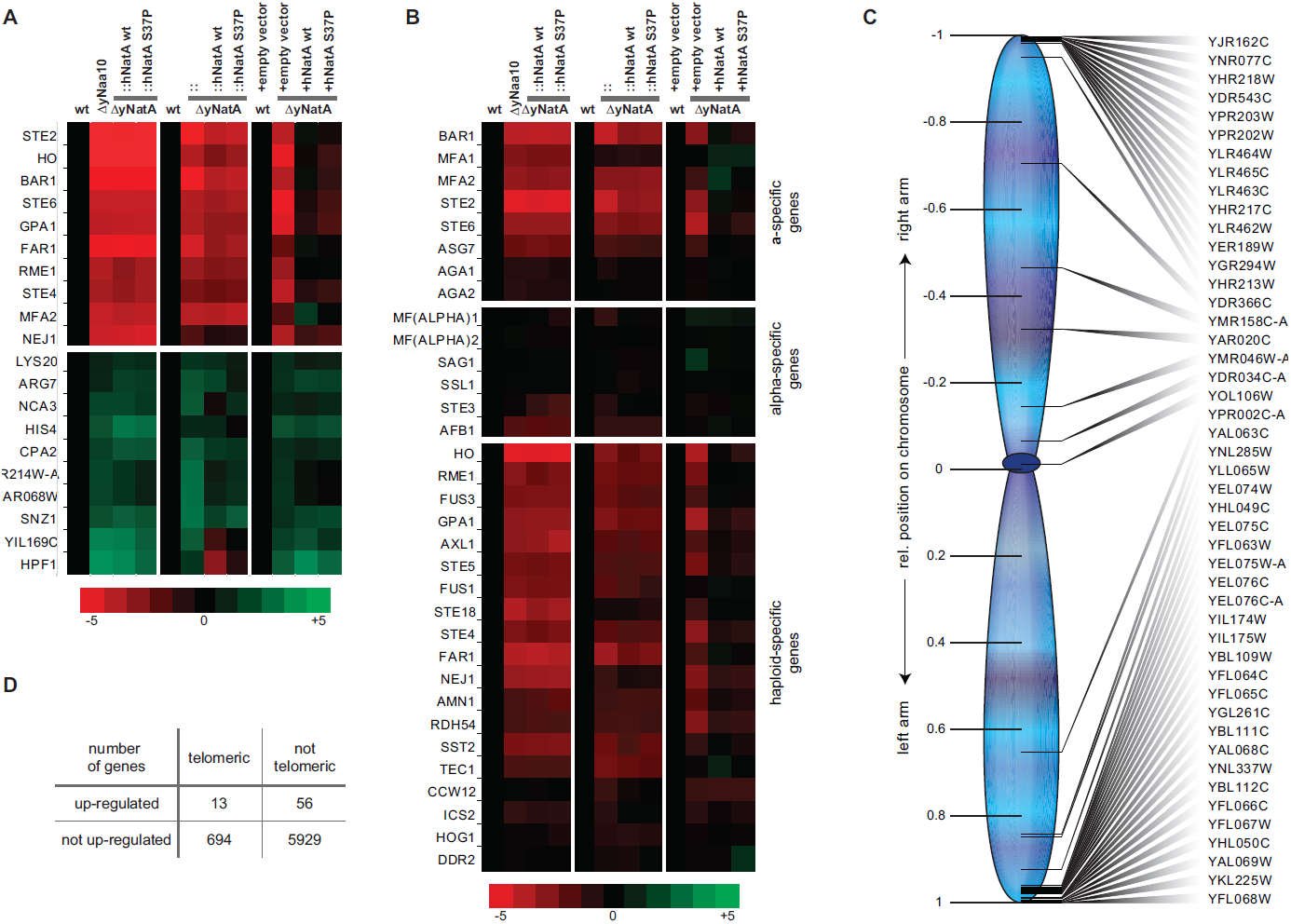
Gene expression analysis. Heat map representation of the microarray results for the the Top 10 up-or downregulated genes in all knockout strains (A) and mating type specific genes (B). C) Presentation of the relative position on the chromosome of the genes that were found to be upregulated by at least 3–fold in the yNatA knockout (overexpressing strains only). D) Telomeric enrichment analysis for the RNA-seq data. A 2–fold expression change and a BH adjusted p-values of less than 0.05 was used to identify up-regulated genes in the knockout compared to WT (69 genes). 40kb was used as the telomeric boundary (707 genes).

Among the genes that are downregulated in the knockout strains of all sets by log2 FC ≤ – 0. 5 (117 genes) a GO term analysis using the WebGestalt revealed enrichments for mating and fusion. The top hits were “cell surface receptor signaling pathway” (GO:0007166, adjP=7.68e“^−07^), “pheromone-dependent signal transduction involved in conjugation with cellular fusion” (GO:0000750, adjP=7.68e“^−07^) and “response to pheromone” (GO:0019236, adjP=7.68e” ^−07^). Similar results were obtained when using the RNA-seq data (not shown). Early studies showed that the depletion of Naa10 results in the de-repression of the normally silent α-information at the *HML* locus ^55; 62^. Therefore, we speculated that the induced expression of the alpha1/2 proteins would radically change the expression profile in ΔNatA*MAT***a** cells. Indeed, a detailed analysis of mating-type specific genes showed a down-regulation of **a**-specific genes such as the receptor for alpha pheromone, Ste2, and haploid-specific genes (e.g. HO) in the yNaa10/NatA knockout cells (Figure 7B).

A down-regulation of α-specific genes in the *MAT***a** cells was not observed. A similar silencing mechanism has been described for the telomeric regions [for a review see ^63^]. Since we expected that a disruption of telomeric silencing would increase the expression of genes specifically in that region, we plotted the relative position of the most up-regulated genes in the knockout according to their relative position on the chromosomes. As shown in Figure 7C for the overexpressing strains set, these genes cluster toward the end of the chromosomes, indeed indicating a de-repression of subtelomeric regions. This effect could also be observed in the endogenous set, albeit to a lesser extent (not shown). Overexpression of the human NatA complex did not restore the repression of these genes in the subtelomeric region (Figure 7C). A similar analysis of telomeric region expressions was also performed using RNA-seq data. By considering log2 fold change > 1 and BH adjusted p-values less than 0.05, 69 genes are found to be up-regulated in the knockout cells, relative to the wild type (Figure 7D). Consistent with microarray data, we found that up-regulated genes are significantly enriched in the telomeric regions (p-value=0.04, Chi-square test). Together, these results suggest that the disruption of telomeric silencing due to knock out of the yeast NatA complex increases the gene expression levels in those regions.

### Knockout of Naa10/NatA affects mating

The pseudo-diploid gene expression pattern in *MAT***a** Naa10/NatA knockout yeast should interfere with the ability of the cells to mate. To test this hypothesis, we performed quantitative mating experiments. Indeed, the knockout of the NatA complex completely abolished mating in yeast (Figure 8). The overexpression of the human NatA complex (hNatA) rescued the mating ability of the cells to WT levels; however, overexpression of the mutated Naa10 S37P/Naa15 complex showed a statistically significant diminished mating when compared to the WT strain.

**Figure 8:**
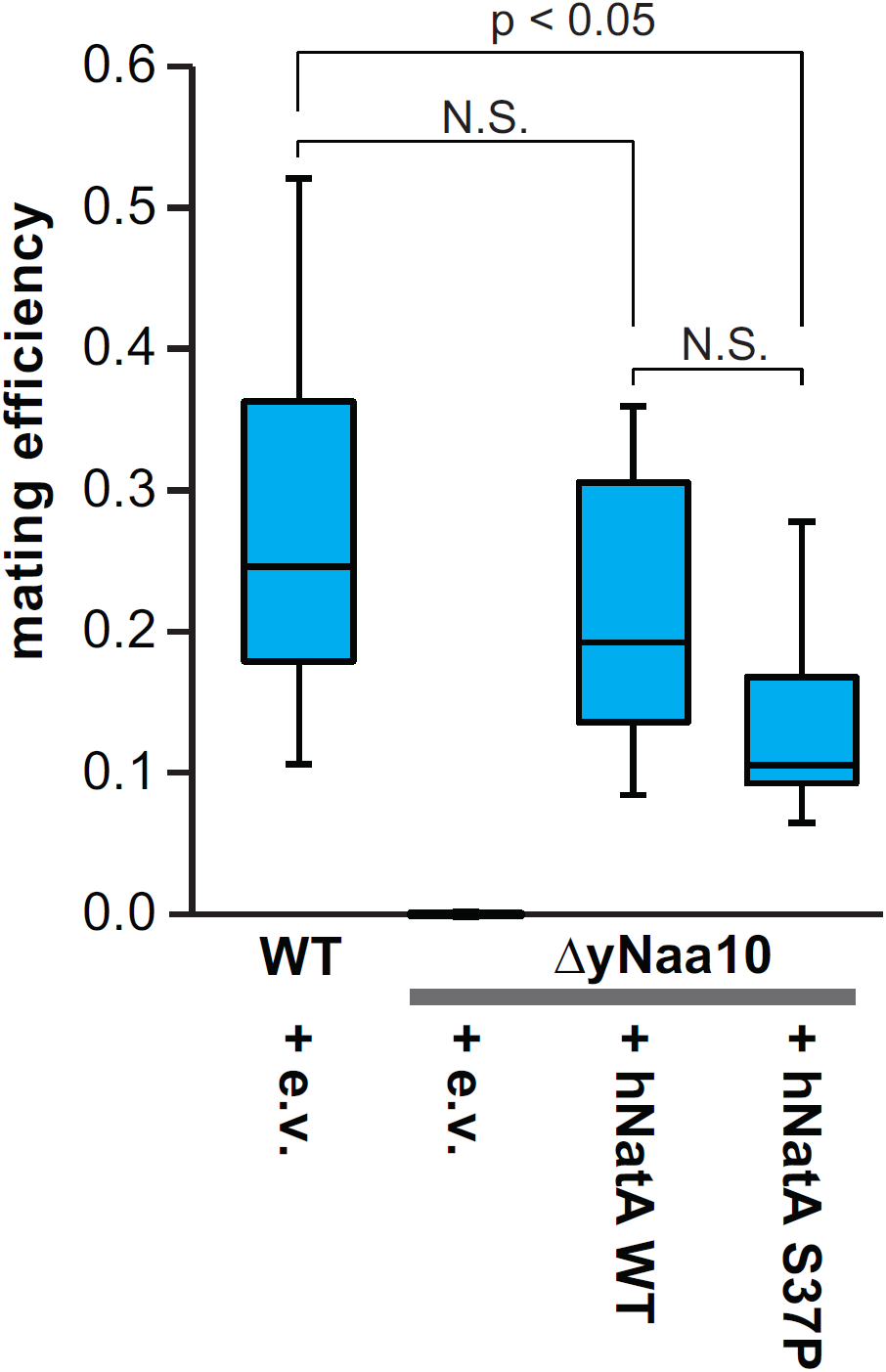
Quantitative mating experiments. To quantify mating efficiencies, cells were seeded on SC“^Lys-Ura^ selective plates containing a lawn of mating tester cells and in parallel on SC^−Ura^ plates to determine the seeding cell number. Cell colonies were counted after 2–3 days and the mating efficiency was determined by calculating the ratio between the mated and the seeded cells. Shown is a box plot representation for the results from 9 independent experiments. Statistical analyses were performed using student's t-test.

## Discussion

Here we show that knockout of the yeast NatA complex induced a strong growth defect at elevated temperatures in liquid culture and on plates. This is in line with previous findings that knockout of Naa10 or Naa15 induces heat sensitivity in yeast ^11; 55; 56; 64^ Interestingly, overexpression of the human NatA complex rescued this growth defect at 39°C whereas the mutated hNaa10 S37P/hNaa15 only partially restored the defect, indicating a functional impairment of the Ogden mutation *in vivo*. The reason for the heat sensitivity is not known to date; however, GO term enrichment analyses from our proteomic characterization suggest that disruption of NatA might perturb protein folding and re-folding, which could contribute to this phenotype. Furthermore, a recent study showed that disruption of NatA in a [PSI+] prion strain led to the accumulation of protein aggregates and the activation of heat-shock response proteins of the Hsp104, Hsp70 and Hsp40 family ^61^. Similarly, in our mass spectrometric analyses, Hsp104, Ssa1–4 (Hsp70) and Sis1 (Hsp40) were increased in the knockout under normal growth conditions, whereas they were back to control levels upon overexpression of hNatA. Overexpressing of the mutated hNatA S37P complex only partially compensated this effect, which is in line with the partial rescue observed in the growth tests. This finding indicates that the observed effects may be caused by an impaired functionality of these chaperones, possibly due to a reduced acetylation of their N-termini. The up-regulation of these proteins could be a naturally occurring mechanism of the cells to compensate for their diminished activity. Indeed, Ssa1-4 and Ssb1/2 proteins have been shown to be NatA substrates ^40; 51^ and loss of NTA of Ssa1/2 and Ssb1/2 has been shown to disturb their functionality in the propagation of the prion phenotype ^61^. Additionally, overexpression of the heat-inducible Hsp70 proteins Ssb1/2 could partially suppress the temperature sensitivity in ΔyNaa15 or ΔyNaa10 cells ^36^. This suggests that loss of NTA of the Hsp70 chaperones Ssa1–4 and Ssb1/2 in ΔNatA cells reduces the functionality of these proteins, thereby affecting protein biogenesis and reducing cellular fitness under normal conditions as seen in the liquid growth experiments. Similarly, primary fibroblasts from the Ogden patient exhibit a decreased growth rate and proliferation when compared to control cells ^45^. Under stress conditions, when there is a heavy load on protein folding, this effect could be further aggravated as seen in the heat shock experiments presented here. It would be interesting to see if an elimination of NTA from Ssa1/4 and Ssb1/2 is sufficient to reproduce the heat sensitivity or if the defect of ANatA cells could be restored to WT levels by artificially acetylating these proteins. The latter could be done in future experiments by mutating the second (first after iMet excision) amino acid of these proteins to change their NAT specificity and/or to prevent their NTA entirely, similarly to the experiments performed for the propagation of the prion phenotype ^61^.

In analogy to the temperature challenge, similar results were obtained when cycloheximide was used as chemical stressor. Knockout of yNatA strongly decreased the resistance of yeast toward cycloheximide and this defect could be partially rescued by overexpression of the human hNat15/hNaa10 complex from plasmids. The hNaa10 S37P mutant showed a similar or slightly decreased rescue effect, when compared to hNaa10 WT. This is in line with previous experiments where the sensitivity of ΔyNatA yeast toward cycloheximide and caffeine was nearly completely repressed by the expression of hNatA, whereas expression of hNatA S37P resulted in a reduced complementation ^11; 51^.

A similar trend could be observed when we used doxorubicin as a stressor; however, in this case Naa10 negatively regulates cell survival. In fact, knockout of yNatA improved survival of the cells, whereas overexpression of the hNatA reduced cell growth on doxorubicin-containing plates. From studies in *D. melanogaster* and HeLa, Ht1080 and U2OS cells, it is known that siRNA-mediated knockdown of Naa10 protects the cells from doxorubicin induced cell death ^65; 66^ which indicates that Naa10 has a regulatory function in DNA damage control. Indeed, doxorubicin treatment induced RIP1/PIDD/NEMO complex formation, NEMO ubiquitination and NF-*κ*B activation in HEK293 ^67^, providing a possible mechanism for Naa10 function in DNA damage. However, the NF-κB pathway is absent in yeast ^68^, so the mechanism in yeast needs further study. Similarly to the effects observed for doxorubicin, knockdown of Naa10 also increased resistance of HeLa cells toward cisplatin and ultraviolet treatment ^66^. In contrast to this, we did not observe any growth or survival advantage of a NatA knockout in yeast under conditions of UV irradiation.

Interestingly, the replacement of the yeast NatA complex with the human genes on the endogenous locus did not reproducibly rescue the heat shock defects seen in the knockout, along with more variability noted among the different biological replicates (Figure 2). One explanation for this finding is that the human proteins might not be expressed as efficiently as the corresponding yeast proteins in *S. cerevisiae* (also see Figure 1). Furthermore, despite the NatA complex being evolutionarily conserved from yeast to vertebrates ^69; 70^, with all tested variants having a similar substrate specificity, slight structural differences could result in the human proteins being less functional in yeast. A comparative N-terminal acetylome analysis using COFRADIC revealed that, although human and yeast NatA acetylate the same set of proteins, hNatA displays a preference toward Ala-N-termini whereas yNatA seems to be the more efficient in acetylating Ser-starting N-termini ^11; 51^. The latter study showed that yNatA seems to be more efficient than the human complex in yeast ^51^. Hence, the expression of the human proteins at the endogenous locus might not be sufficient to complement yeast NatA, whereas overexpression from a plasmid could make up for a diminished acetylation activity of the human proteins toward yeast substrates. Similarly, overexpression of the human proteins rather than endogenous expression was necessary to revert the expression changes induced by disruption of yNatA. Interestingly, endogenous expression of the human proteins partially rescued a growth defect toward cycloheximide treatment, which could indicate that NatA substrates involved in the cycloheximide response are less susceptible to a reduction in NTA.

In contrast to the effects observed with the hNatA S37P mutation, the homologous mutation in yeast Naa10 showed no observable effects. This suggests that there are significant structural differences between the species. In this regard it has been shown that heterologous combinations of human and yeast Naa10/Naa15 are not functional in yeast ^11^. Also, the sequence alignment performed in this study showed an additional putative α-helix C-terminally adjacent to the S39 in *S. cerevisiae* which is missing in the human Naa10. Furthermore, the Ogden mutation is not completely deleterious in human Naa10 since residual catalytic activity of Naa10 S37P could be found *in vitro* ^44^ in a yeast model ^51^ and in primary cells from the patients ^45^. Therefore, structural difference and the relatively mild nature of the mutation itself could explain the absence of an obvert phenotype in yeast.

Taken together, the data presented here indicates that Naa10 positively regulates cell survival in response to cycloheximide and heat stress but promotes cell death in response to doxorubicin. The human NatA proteins could rescue the yeast knockout, at least when overexpressed, with the Naa10 S37P mutation being less efficient in most experiments, indicating that the Ogden mutation is affecting Naa10 function *in vivo*, supporting the disruptive nature of this mutation.

To analyze the effects of NatA disruption at the transcriptional level, we performed microarray studies on 3 different sets, each containing a WT strain, a Naa10/NatA knockout and 2 rescue strains expressing the human NatA WT or S37P proteins, respectively. A GO-term enrichment analysis of the down-regulated genes in the knockout revealed that these genes were involved in mating, cellular fusion and response to pheromone. We also found a similar enrichment in the mass spectrometric analyses, highlighting that the down-regulation of genes involved in mating/fusion can also be detected at the protein level.

In *S. cerevisiae*, mating is regulated by **a**-and α-specific transcription factors that are encoded by the nonhomologous alleles *MAT**a*** and*MAT*α^71^. The ***MAT*** locus is encompassed by two silenced alleles, *HML* and *HMR*, that contains a cryptic copy of the mating type sequences *MAT***a** and *MAT*α, respectively, that serve as donors for the *MAT* locus to confer the mating type of a haploid cell ^71^. Silencing of the *HML* and *HMR* is accomplished by trans-acting factors that bind to specific silencer sequences surrounding these loci. First, the origin recognition complex ORC binds to the silencer and recruits silent information regulator, Sir1^72^. This recruits the Sir2/Sir3/Sir4 complex which de-acetylates the tails of histone H3 and H4, thereby creating additional binding sites for Sir3/Sir4, resulting in a sequential spreading of these proteins and transcriptional repression in this region^73^. Sir3 and Orc1, the large subunit of the origin recognition complex, are both substrates of NatA, and NTA of these proteins stabilizes their binding to nucleosomes, thereby facilitating silencing of *HML*/*HMRL*^74–77^. Disruption of NatA should therefore induce the expression of **a**-and α-genes from the de-repressed *HML*/*HMR* loci, leading to a pseudo-diploid gene expression profile. Our microarray and RNA-seq data supports this hypothesis: **a**-and haploid-specific genes were down-regulated in ΔyNaa10/yNatA knockout cells, when compared to the WT. In haploid **a**-cells, as used in this study, alpha-specific genes are constitutively repressed. Diploid cells contain both, *MAT***a** and *MAT*α, which results in the formation of the **a**/alpha2 repressor that keeps alpha-specific genes repressed^78^. In analogy to this, the artificial expression of *HML*/*HMR* genes as a result of the yNaa10/yNatA knockout did not affect the gene expression of alpha-specific genes in our analysis, instead they remained repressed. This pseudo-diploid expression profile (e.g. down-regulation of the receptor for alpha-factor, Ste2 or repression of Ste4, a G protein involved in the mating signaling pathway) was accompanied by a dramatically reduced mating efficiency as shown in quantitative mating experiments. This is in line with previous findings that disruption of NatA interferes with mating^55; 62; 79^. This mating effect is seen only in *MAT***a** cells, as used in this study, since silencing at *HML* is less stable than at *HMR*, especially when Sir3 becomes limiting ^76; 80^.

In agreement with the stress tests, the pseudo-diploid gene expression pattern as well as the mating efficiency could be partially reversed by overexpression of the human NatA complex. Furthermore, overexpression of the Ogden mutant was less efficient in rescuing the mating defect.

Sir3 is also involved in silencing at telomeres^76; 81^. An impaired functionality of this protein as a result of their reduced NTA should therefore lead to de-repression of genes in that region. Consistent with this, we could detect an up-regulation of genes in sub-telomeric regions in the NatA knockout strains by RNA-seq and microarray. Also, in contrast to the pseudo-diploid expression profile, the de-repression of genes at telomeres could not be complemented by overexpression of the human NatA proteins. It is known that telomeric silencing is less robust than *HML*/*HMR* silencing ^71^ and might therefore be more susceptible to small perturbations. Since the human NatA proteins seemed to be less functional toward yeast proteins (see above), it could be possible that the overexpression of hNatA only partially restored NTA of Sir3 and Orc1, which in turn might be sufficient to restore *HML*/*HMR* but not telomeric silencing.

In conclusion, we have shown here that disruption of NatA leads to growth defects and increased sensitivity toward stresses such as heat or cycloheximide in *S. cerevisiae*. These effects are possibly due to an impaired functionality of molecular chaperones. Additionally, ΔNatA cells are characterized by a pseudo-diploid gene expression pattern and failure to mate, probably due to reduced NTA of the NatA substrates Orc1 and Sir3. These effects could be restored by expression of the human NatA complex to different degrees, depending on the system used (endogenous expression vs. overexpression) and other cellular processes. Interestingly, a mutated form of hNaa10, harboring the variant implicated in Ogden syndrome, displayed a similar or diminished complementation when compared to WT hNaa10. This strongly supports the disruptive nature of the Ogden/S37P mutation in Naa10, consistent with prior results^45; 51^.

## Acknowledgements

Sabine Rospert (Germany) provided anti-yeast Naa10 and Naa15 antibodies. The W303-1A WT strain (YG2) as well as the deletion strains ΔyNaa10 (YG1), ΔyNaa10 (YG10) and ΔyNaa15 (YG3) were provided by Rolf Sternglanz. The overexpression set (strain expressing set with the human proteins from plasmids, YG32–35) was provided by Thomas Arnesen. The mating tester strains (YG62, YG63) were kindly provided by Aaron Neiman (USA). We thank Maitreya Dunham (USA) and her team for performing the RNA microarrays. We acknowledge the assistance of Keith Rivera and Darryl Pappin in the CSHL proteomics core facility. Jason O’Rawe in the Lyon lab provided bioinformatics and statistical analysis support.

## Author contributions

MJD planned and performed experiments, analyzed data, prepared all figures, drafted and revised the manuscript. JC provided technical assistance. HF analyzed the RNA-seq data. MK generated yeast strains and commented on the manuscript. JW performed the experiments for figure 8. GJL planned the study, provided experimental guidance, and revised the manuscript.

## References

1. Deakin,J.F., Döstrovsky,J.O., and Smyth,D.G. 1980. Influence of N-terminal acetylation and C-terminal proteolysis on the analgesic activity of beta-endorphin. Biochem J 189, 501–506.

2. Scott,D.C., Monda,J.K., Bennett,E.J., Harper,J.W., and Schulman,B.A. 2011. N-terminal acetylation acts as an avidity enhancer within an interconnected multiprotein complex. Science 334, 674–678.

3. Nazmi,A.R., Ozorowski,G., Pejic,M., Whitelegge,J.P., Gerke,V., and Luecke,H. 2012. N-terminal acetylation of annexin A2 is required for S100A10 binding. Biol Chem 393, 1141–1150.

4. Hwang,C.S., Shemorry,A., and Varshavsky,A. 2010. N-terminal acetylation of cellular proteins creates specific degradation signals. Science 327, 973–977.

5. Shemorry,A., Hwang,C.S, and Varshavsky,A. 2013. Control of protein quality and stoichiometries by N-terminal acetylation and the N-end rule pathway. Mol Cell 50, 540–551.

6. Park,S.E., Kim,J.M, Seok,O.H, Cho,H., Wadas,B., Kim,S.Y, Varshavsky,A., and Hwang,C. S. 2015. Control of mammalian G protein signaling by N-terminal acetylation and the N-end rule pathway. Science 347, 1249–1252.

7. Ben-Saadon,R., Fajerman,I., Ziv,T., Hellman,U., Schwartz,A.L, and Ciechanover,A. 2004. The tumor suppressor protein p16(INK4a) and the human papillomavirus oncoprotein-58 E7 are naturally occurring lysine-less proteins that are degraded by the ubiquitin system. Direct evidence for ubiquitination at the N-terminal residue. J Biol Chem 279, 41414–41421.

8. Ciechanover,A., and Ben-Saadon,R. 2004. N-terminal ubiquitination: more protein substrates join in. Trends Cell Biol 14, 103–106.

9. Kuo,M.L., den Besten,W., and Sherr,C.J 2004. N-Terminal polyubiquitination of the ARF tumor suppressor, a natural lysine-less protein. Cell Cycle 3, 1367–1369.

10. Aksnes,H., Hole,K., and Arnesen,T. 2015. Molecular, cellular, and physiological significance of N-terminal acetylation. Int Rev Cell Mol Biol 316, 267–305.

11. Arnesen,T., Van Damme,P., Polevoda,B., Helsens,K., Evjenth,R., Colaert,N., Varhaug,J.E., Vandekerckhove,J., Lillehaug,J.R, Sherman,F., et al. 2009. Proteomics analyses reveal the evolutionary conservation and divergence of N-terminal acetyltransferases from yeast and humans. Proc Natl Acad Sci U S A 106, 8157–8162.

12. Van Damme,P., Evjenth,R., Foyn,H., Demeyer,K., De Bock,P.J., Lillehaug,J.R, Vandekerckhove,J., Arnesen,T., and Gevaert,K. 2011. Proteome-derived peptide libraries allow detailed analysis of the substrate specificities of N(alpha)-acetyltransferases and point to hNaa10p as the post-translational actin N(alpha)-acetyltransferase. Mol Cell Proteomics 10, M110.004580.

13. Lange,P.F., Huesgen,P.F, Nguyen,K., and Overall,C.M 2014. Annotating N termini for the human proteome project: N termini and Nα-acetylation status differentiate stable cleaved protein species from degradation remnants in the human erythrocyte proteome. J Proteome Res 13, 2028–2044.

14. Shoemaker,K.R., Kim,P.S, York,E.J, Stewart,J.M, and Baldwin,R.L 1987. Tests of the helix dipole model for stabilization of alpha-helices. Nature 326, 563–567.

15. Fairman,R., Shoemaker,K.R, York,E.J, Stewart,J.M, and Baldwin,R.L 1989. Further studies of the helix dipole model: effects of a free alpha-NH3+ or alpha-COO-group on helix stability. Proteins 5, 1–7.

16. Doig,A.J., Chakrabartty,A., Klingler,T.M, and Baldwin,R.L 1994. Determination of free energies of N-capping in alpha-helices by modification of the Lifson-Roig helix-coil therapy to include N-and C-capping. Biochemistry 33, 3396–3403.

17. Greenfield,N.J., Stafford,W.F, and Hitchcock-DeGregori,S.E. 1994. The effect of N-terminal acetylation on the structure of an N-terminal tropomyosin peptide and alpha alpha-tropomyosin. Protein Sci 3, 402–410.

18. Jarvis,J.A., Ryan,M.T, Hoogenraad,N.J, Craik,D.J, and Høj,P.B. 1995. Solution structure of the acetylated and noncleavable mitochondrial targeting signal of rat chaperonin 10. J Biol Chem 270, 1323–1331.

19. Fauvet,B., Fares,M.B, Samuel,F., Dikiy,I., Tandon,A., Eliezer,D., and Lashuel,H.A (2012). Characterization of Semisynthetic and Naturally Nα-Acetylated α-Synuclein in Vitro and in Intact Cells: IMPLICATIONS FOR AGGREGATION AND CELLULAR PROPERTIES OF α-SYNUCLEIN. J Biol Chem 287, 28243–28262.

20. Kang,L., Moriarty,G.M, Woods,L.A, Ashcroft,A.E, Radford,S.E, and Baum,J. 2012. N-terminal acetylation of α-synuclein induces increased transient helical propensity and decreased aggregation rates in the intrinsically disordered monomer. Protein Sci 21, 911–917.

21. Kang,L., Janowska,M.K, Moriarty,G.M, and Baum,J. 2013. Mechanistic Insight into the Relationship between N-Terminal Acetylation of α-Synuclein and Fibril Formation Rates by NMR and Fluorescence. PLoS One 8, e75018.

22. Permyakov,S.E., Vologzhannikova,A.A, Emelyanenko,V.I, Knyazeva,E.L, Kazakov,A.S., Lapteva,Y.S, Permyakova,M.E, Zhadan,A.P, and Permyakov,E.A 2012. The impact of alpha-N-acetylation on structural and functional status of parvalbumin. Cell Calcium 52, 366–376.

23. Scheepens,A., Mould,R., Hofmann,O., and Brittain,T. 1995. Some effects of post-translational N-terminal acetylation of the human embryonic zeta globin protein. Biochem J 310 ( Pt 2), 597–600.

24. Ashiuchi,M., Yagami,T., Willey,R.J, Padovan,J.C, Chait,B.T, Popowicz,A., Manning,L.R., and Manning,J.M 2005. N-terminal acetylation and protonation of individual hemoglobin subunits: position-dependent effects on tetramer strength and cooperativity. Protein Sci 14, 1458–1471.

25. Starling,A.P., Sharma,R.P, East,J.M, and Lee,A.G 1996. The effect of N-terminal acetylation on Ca(2+)-ATPase inhibition by phospholamban. Biochem Biophys Res Commun 226, 352–355.

26. Van Doren,S.R, Wei,S., Gao,G., DaGue,B.B, Palmier,M.O, Bahudhanapati,H., and Brew,K. 2008. Inactivation of N-TIMP-1 by N-terminal acetylation when expressed in bacteria. Biopolymers 89, 960–968.

27. Polevoda,B., Cardillo,T.S, Doyle,T.C, Bedi,G.S, and Sherman,F. 2003. Nat3p and Mdm20p are required for function of yeast NatB Nalpha-terminal acetyltransferase and of actin and tropomyosin. J Biol Chem 278, 30686–30697.

28. Singer,J.M., and Shaw,J.M 2003. Mdm20 protein functions with Nat3 protein to acetylate Tpm1 protein and regulate tropomyosin-actin interactions in budding yeast. Proc Natl Acad Sci U S A 100, 7644–7649.

29. Coulton,A.T., East,D.A, Galinska-Rakoczy,A., Lehman,W., and Mulvihill,D.P 2010. The recruitment of acetylated and unacetylated tropomyosin to distinct actin polymers permits the discrete regulation of specific myosins in fission yeast. J Cell Sci 123, 3235–3243.

30. Caesar,R., and Blomberg,A. 2004. The stress-induced Tfs1p requires NatB-mediated acetylation to inhibit carboxypeptidase Y and to regulate the protein kinase A pathway. J Biol Chem 279, 38532–38543.

31. Kalvik,T.V., and Arnesen,T. 2013. Protein N-terminal acetyltransferases in cancer. Oncogene 32, 269–276.

32. Dörfel,M.J., and Lyon,G.J 2015. The biological functions of Naa10-From amino-terminal acetylation to human disease. Gene 567, 103–131.

33. Polevoda,B., Arnesen,T., and Sherman,F. 2009. A synopsis of eukaryotic Nalpha-terminal acetyltransferases: nomenclature, subunits and substrates. BMC Proc 3 Suppl 6, S2.

34. Van Damme,P., Hole,K., Pimenta-Marques,A., Helsens,K., Vandekerckhove,J., Martinho,R.G., Gevaert,K., and Arnesen,T. 2011. NatF contributes to an evolutionary shift in protein N-terminal acetylation and is important for normal chromosome segregation. PLoS Genet 7, e1002169.

35. Starheim,K.K., Gevaert,K., and Arnesen,T. 2012. Protein N-terminal acetyltransferases: when the start matters. Trends Biochem Sci 37, 152–161.

36. Gautschi,M., Just,S., Mun,A., Ross,S., Rücknagel,P., Dubaquié,Y., Ehrenhofer-Murray,A., and Rospert,S. 2003. The yeast N(alpha)-acetyltransferase NatA is quantitatively anchored to the ribosome and interacts with nascent polypeptides. Mol Cell Biol 23, 7403–7414.

37. Raue,U., Oellerer,S., and Rospert,S. 2007. Association of prot in biogenesis factors at the yeast ribosomal tunnel exit is affected by the translational status and nascent polypeptide sequence. J Biol Chem 282, 7809–7816.

38. Arnesen,T., Gromyko,D., Kagabo,D., Betts,M.J, Starheim,K.K, Varhaug,J.E, Anderson,D., and Lillehaug,J.R 2009. A novel human NatA Nalpha-terminal acetyltransferase complex: hNaa16p-hNaa10p (hNat2-hArd1). BMC Biochem 10, 15.

39. Arnold,R.J., Polevoda,B., Reilly,J.P, and Sherman,F. 1999. The action of N-terminal acetyltransferases on yeast ribosomal proteins. J Biol Chem 274, 37035–37040.

40. Polevoda,B., Norbeck,J., Takakura,H., Blomberg,A., and Sherman,F. 1999. Identification and specificities of N-terminal acetyltransferases from Saccharomyces cerevisiae. EMBO J 18, 6155–6168.

41. Foyn,H., Jones,J.E, Lewallen,D., Narawane,R., Varhaug,J.E, Thompson,P.R, and Arnesen,T. 2013. Design, Synthesis, and Kinetic Characterization of Protein N-Terminal Acetyltransferase Inhibitors. ACS Chem Biol.

42. Liszczak,G., Goldberg,J.M, Foyn,H., Petersson,E.J, Arnesen,T., and Marmorstein,R. (2013). Molecular basis for N-terminal acetylation by the heterodimeric NatA complex. Nat Struct Mol Biol.

43. Magin,R.S., March,Z.M, and Marmorstein,R. 2016. The N-terminal acetyltransferase Naa10/ARD1 does not acetylate lysine residues. J Biol Chem.

44. Rope,A.F., Wang,K., Evjenth,R., Xing,J., Johnston,J.J, Swensen,J.J, Johnson,W.E, Moore,B., Huff,C.D, Bird,L.M, et al. 2011. Using VAAST to identify an X-linked disorder resulting in lethality in male infants due to N-terminal acetyltransferase deficiency. Am J Hum Genet 89, 28–43.

45. Myklebust,L.M., Van Damme,P., Stove,S.I, Dorfel,M.J, Abboud,A., Kalvik,T.V, Grauffel,C., Jonckheere,V., Wu,Y., Swensen,J., et al. 2015. Biochemical and cellular analysis of Ogden syndrome reveals downstream Nt-acetylation defects. Human molecular genetics 24, 1956–1976.

46. Van Damme,P., Støve,S.I., Glomnes,N., Gevaert,K., and Arnesen,T. 2014. A Saccharomyces cerevisiae model reveals in vivo functional impairment of the Ogden syndrome N-terminal acetyltransferase Naa10S37P mutant. Mol Cell Proteomics.

47. Rauch,A., Wieczorek,D., Graf,E., Wieland,T., Endele,S., Schwarzmayr,T., Albrecht,B.,Bartholdi,D., Beygo,J., Di Donato,N., et al. 2012. Range of genetic mutations associated with severe non-syndromic sporadic intellectual disability: an exome sequencing study. Lancet 380, 1674–1682.

48. Popp,B., Støve,S.I., Endele,S., Myklebust,L.M, Hoyer,J., Sticht,H., Azzarello-Burri,S.,Rauch,A., Arnesen,T., and Reis,A. 2014. De novo missense mutations in the NAA10 gene cause severe non-syndromic developmental delay in males and females. Eur J Hum Genet.

49. Casey,J.P., Støve,S.I., McGorrian,C., Galvin,J., Blenski,M., Dunne,A., Ennis,S., Brett,F., King,M.D., Arnesen,T., et al. 2015. NAA10 mutation causing a novel intellectual disability syndrome with Long QT due to N-terminal acetyltransferase impairment. Sci Rep 5, 16022.

50. Esmailpour,T., Riazifar,H., Liu,L., Donkervoort,S., Huang,V.H, Madaan,S., Shoucri,B. M., Busch,A., Wu,J., Towbin,A., et al. 2014. A splice donor mutation in NAA10 results in the dysregulation of the retinoic acid signalling pathway and causes Lenz microphthalmia syndrome. J Med Genet.

51. Van Damme,P., Støve,S.I., Glomnes,N., Gevaert,K., and Arnesen,T. 2014. A Saccharomyces cerevisiae model reveals in vivo functional impairment of the Ogden syndrome N-terminal acetyltransferase NAA10 Ser37Pro mutant. Mol Cell Proteomics 13, 2031–2041.

52. Amberg,D.C., Burke,D., and Strathern,J.N 2005. Methods in Yeast Genetics : A Cold Spring Harbor Laboratory Course Manual. Cold Spring Harbor Laboratory Press.

53. Zilio,N., Wehrkamp-Richter,S., and Boddy,M.N 2012. A new versatile system for rapid control of gene expression in the fission yeast Schizosaccharomyces pombe. Yeast 29, 425–434.

54. Ross,P.L., Huang,Y.N, Marchese,J.N, Williamson,B., Parker,K., Hattan,S., Khainovski,N., Pillai,S., Dey,S., Daniels,S., et al. 2004. Multiplexed protein quantitation in Saccharomyces cerevisiae using amine-reactive isobaric tagging reagents. Mol Cell Proteomics 3, 1154–1169.

55. Whiteway,M., and Szostak,J.W 1985. The ARD1 gene of yeast functions in the switch between the mitotic cell cycle and alternative developmental pathways. Cell 43, 483–492.

56. Mullen,J.R., Kayne,P.S, Moerschell,R.P, Tsunasawa,S., Gribskov,M., Colavito-Shepanski,M., Grunstein,M., Sherman,F., and Sternglanz,R. 1989. Identification and characterization of genes and mutants for an N-terminal acetyltransferase from yeast. EMBO J 8, 2067–2075.

57. Lee,F.J., Lin,L.W, and Smith,J.A 1989. N alpha acetylation is required for normal growth and mating of Saccharomyces cerevisiae. J Bacteriol 171, 5795–5802.

58. Patel,S., Sprung,A.U, Keller,B.A, Heaton,V.J, and Fisher,L.M 1997. Identification of yeast DNA topoisomerase II mutants resistant to the antitumor drug doxorubicin: implications for the mechanisms of doxorubicin action and cytotoxicity. Molecular pharmacology 52, 658–666.

59. Wang,J., Duncan,D., Shi,Z., and Zhang,B. 2013. WEB-based GEne SeT AnaLysis Toolkit (WebGestalt): update 2013. Nucleic Acids Res 41, W77–83.

60. Zhang,B., Kirov,S., and Snoddy,J. 2005. WebGestalt: an integrated system for exploring gene sets in various biological contexts. Nucleic Acids Res 33, W741–748.

61. Holmes,W.M., Mannakee,B.K, Gutenkunst,R.N, and Serio,T.R 2014. Loss of amino-terminal acetylation suppresses a prion phenotype by modulating global protein folding. Nat Commun 5, 4383.

62. Whiteway,M., Freedman,R., Van Arsdell,S., Szostak,J.W, and Thorner,J. 1987. The yeast ARD1 gene product is required for repression of cryptic mating-type information at the HML locus. Mol Cell Biol 7, 3713–3722.

63. Gartenberg,M.R. 2000. The Sir proteins of Saccharomyces cerevisiae: mediators of transcriptional silencing and much more. Current opinion in microbiology 3, 132–137.

64. Lee,F.J., Lin,L.W, and Smith,J.A 1989. N alpha-acetyltransferase deficiency alters protein synthesis in Saccharomyces cerevisiae. FEBS Lett 256, 139–142.

65. Yi,C.H., Sogah,D.K, Boyce,M., Degterev,A., Christofferson,D.E, and Yuan,J. 2007. A genome-wide RNAi screen reveals multiple regulators of caspase activation. J Cell Biol 179, 619–626.

66. Yi,C.H., Pan,H., Seebacher,J., Jang,I.H, Hyberts,S.G, Heffron,G.J, Vander Heiden,M.G, Yang,R., Li,F., Locasale,J.W, et al. 2011. Metabolic regulation of protein N-alpha-acetylation by Bcl-xL promotes cell survival. Cell 146, 607–620.

67. Park,J., Kanayama,A., Yamamoto,K., and Miyamoto,Y. 2012. ARD1 binding to RIP1 mediates doxorubicin-induced NF-κB activation. Biochem Biophys Res Commun 422, 291–297.

68. Srinivasan,V., Kriete,A., Sacan,A., and Jazwinski,S.M 2010. Comparing the yeast retrograde response and NF-kappaB stress responses: implications for aging. Aging cell 9, 933–941.

69. Sugiura,N., Adams,S.M, and Corriveau,R.A 2003. An evolutionarily conserved N-terminal acetyltransferase complex associated with neuronal development. J Biol Chem 278, 40113–40120.

70. Arnesen,T., Anderson,D., Baldersheim,C., Lanotte,M., Varhaug,J.E, and Lillehaug,J.R (2005). Identification and characterization of the human ARD1-NATH protein acetyltransferase complex. Biochem J 386, 433–443.

71. Haber,J.E. 2012. Mating-type genes and MAT switching in Saccharomyces cerevisiae. Genetics 191, 33–64.

72. Fox,C.A., Ehrenhofer-Murray,A.E., Loo,S., and Rine,J. 1997. The origin recognition complex, SIR1, and the S phase requirement for silencing. Science 276, 1547–1551.

73. Rusche,L.N., Kirchmaier,A.L, and Rine,J. 2003. The establishment, inheritance, and function of silenced chromatin in Saccharomyces cerevisiae. Annu Rev Biochem 72, 481–516.

74. Wang,X., Connelly,J.J, Wang,C.L, and Sternglanz,R. 2004. Importance of the Sir3 N terminus and its acetylation for yeast transcriptional silencing. Genetics 168, 547–551.

75. Arnaudo,N., Fernández,I.S, McLaughlin,S.H, Peak-Chew,S.Y., Rhodes,D., and Martino,F. 2013. The N-terminal acetylation of Sir3 stabilizes its binding to the nucleosome core particle. Nat Struct Mol Biol.

76. Geissenhöner,A., Weise,C., and Ehrenhofer-Murray,A.E. 2004. Dependence of ORC silencing function on NatA-mediated Nalpha acetylation in Saccharomyces cerevisiae. Mol Cell Biol 24, 10300–10312.

77. van Welsem,T., Frederiks,F., Verzijlbergen,K.F, Faber,A.W, Nelson,Z.W, Egan,D.A, Gottschling,D.E., and van Leeuwen,F. 2008. Synthetic lethal screens identify gene silencing processes in yeast and implicate the acetylated amino terminus of Sir3 in recognition of the nucleosome core. Mol Cell Biol 28, 3861–3872.

78. Goutte,C., and Johnson,A.D 1988. a1 protein alters the DNA binding specificity of alpha 2 repressor. Cell 52, 875–882.

79. Aparicio,O.M., Billington,B.L, and Gottschling,D.E 1991. Modifiers of position effect are shared between telomeric and silent mating-type loci in S. cerevisiae. Cell 66, 1279–1287.

80. Motwani,T., Poddar,M., and Holmes,S.G 2012. Sir3 and epigenetic inheritance of silent chromatin in Saccharomyces cerevisiae. Mol Cell Biol 32, 2784–2793.

81. Ruault,M., De Meyer,A., Loïodice,I., and Taddei,A. 2011. Clustering heterochromatin: Sir3 promotes telomere clustering independently of silencing in yeast. J Cell Biol 192, 417–431.

82. Pei,J., Kim,B.H, and Grishin,N.V 2008. PROMALS3D: a tool for multiple protein sequence and structure alignments. Nucleic acids research 36, 2295–2300.

